# *Ankfn1* vestibular defects in zebrafish require mutations in both ancestral and derived paralogs

**DOI:** 10.1101/2021.09.11.459924

**Authors:** Kevin D. Ross, Jie Ren, Ruilin Zhang, Neil C. Chi, Bruce A. Hamilton

## Abstract

How and to what degree gene duplication events create regulatory innovation, redundancy, or neofunctionalization remain important questions in animal evolution and comparative genetics. *Ankfn1* genes are single copy in most invertebrates, partially duplicated in jawed vertebrates, and only the derived copy retained in most mammals. Null mutations in the single mouse homolog have vestibular and neurological abnormalities. Null mutation of the single Drosophila homolog is typically lethal with severe sensorimotor deficits in rare survivors. The functions and potential redundancy of paralogs in species with two copies is not known. Here we define a vestibular role for *Ankfn1* homologs in zebrafish based on simultaneous disruption of each locus. Zebrafish with both paralogs disrupted showed vestibular defects and early lethality from swim bladder inflation failure. One intact copy at either locus was sufficient to prevent major phenotypes. Our results show that vertebrate *Ankfn1* genes are required for vestibular-related functions, with at least partial redundancy between ancestral and derived paralogs.

## Introduction

*Ankfn1* is a recently-annotated gene with an interesting genealogy [1]. Orthologous genes are recognized by the eponymous ankyrin (ANK) and fibronectin type-III (FN3) motifs and three highly conserved non-motif domains. Orthologs are found in all animal lineages studied to date except urochordates and in some sister groups to animals, including choanoflagellates and filastereans. Some unicellular examples encode an aminoterminal CRIB domain. Some unicellular and nearly all invertebrate homologs include a carboxyterminal Ras association (RA) domain. The orthology group shows a single member per genome outside of vertebrates. In an ancestor to jawed vertebrate, the ancestral gene was incompletely duplicated, with the derived paralog losing the RA domain. In an ancestor to therian mammals, the ancestral copy was lost. All animal genomes analyzed to date thus have zero (urochordates), one ancestral (all other invertebrates), one derived (therian mammals), or two (other vertebrates) *Ankfn1* genes.

*Ankfn1* mutations were identified independently by three groups using forward genetics in flies and mice. Transposon insertion mutations in *Drosophila* (*wake* alleles) were recovered in a screen for sleep-related phenotypes [2]. Although originally reported as null, these appear to have been isoform-specific alleles, which prevented expression in some tissues. An RNAi screen to identify genes required for asymmetric cell division in *Drosophila* sensory organ precursor cells identified the same gene as an essential regulator of Numb segregation during cell division; null mutations (*Banderoula* alleles) had severe developmental consequences and poor viability [3]. Banderoula protein interacted physically with Discs-large, and inhibition of both genes elicited tumorigenesis in the *Drosophila* brain. We identified a presumptive null allele of the mouse homolog by positional cloning of a neurological mutation (*nmf9*) marked by vestibular and neurological phenotypes [1]. Subsequent alleles made by genome editing confirmed gene identification by non-complementation and showed the functional importance of a non-motif domain including a highly conserved GLYLGYLK peptide sequence, with even a glycine-to-alanine substitution failing to complement the original null. *Nmf9* mice were viable and fertile, but had several neurological abnormalities–including defects in circadian onset, fear learning, and vestibular function. We also showed that in *Drosophila* frame-shift mutations in distinct domains all had *Banderoula*-like phenotypes. The few flies that survived to adulthood had profound sensorimotor deficits and died prematurely. Flies heterozygous for null mutations had abnormal sleep patterns, broadly consistent with *wake* mutants and demonstrating dosage sensitivity for some phenotypes.

Differences between flies and mice both in domain architecture and in phenotypic severity raise questions about essentiality and conservation of *Ankfn1* function in other animals, especially vertebrate groups that have two paralogous copies. This difference between mammals and other vertebrates is relatively common: ∼15% of human genes are single-copy in humans and have more than one identified homolog in the zebrafish *Danio rerio* [4]. Paralogous genes can have variable functional outcomes. Full redundancy can result in one copy decaying into a pseudogene that has lost function relative to the ancestral gene [5]. Duplication can also lead to new or divergent functions, through neofunctionalization or subfunctionalization [6]. The facile genetics of *D. rerio* makes it an ideal system to test *Ankfn1* paralogous gene function in the typical vertebrate arrangement of one ancestral and one derived copy. The ancestral copy with an intact RA domain is on chromosome 24, while the derived copy without RA is on chromosome 12. Assessing mutations of each homolog independently and in combination should allow direct tests of essentiality, functional redundancy, and possible neofunctionalization after duplication [7]

Here we provide new resolution on the evolution and constraint of the *Ankfn1* gene genealogy and use CRISPR/Cas9 to generate mutations in both *D. rerio* homologs in order to define major functions and test the degree of genetic redundancy between paralogs in a non-mammal vertebrate. We generated a frameshifting deletion allele of the derived chromosome 12 paralog (*Ankfn1*), and both a frameshifting insertion allele and an in-frame deletion allele at the GLYLGYLK site of the ancestral chromosome 24 paralog (*Ankfn1-like*). We observed mutant progeny among digenic crosses and identified an overt swim bladder phenotype that was highly penetrant in fish with biallelic inactivation at both loci, weakly penetrant in fish with only the derived copy inactivated, and indistinguishable from background in fish with only the ancestral copy inactivated. Affected fish did not inflate their swim bladder by 5 days post-fertilization, had severe locomotor and balance deficits, and died prior to adulthood. These results suggest genetic overlap between the ancestral and derived homologs for essential functions, including development of the vestibular system.

## Results

### A highly constrained peptide is a predicted structural element of *Ankfn1* proteins

Previous analysis of 14 diverse metazoan *Ankfn1* homologs identified strong conservation of ANK, FN3, and three non-motif domains based on dN/dS analysis across sliding windows [1]. To better understand conserved features in Ankfn1 proteins, we curated additional homologs, many of which were unavailable or poorly annotated at the time of the earlier analysis (Supplementary Table S1). Surprisingly, annotation of two monotreme species (*Ornithorhynchus anatinus* and *Tachyglossus aculeatus*) include ancestral paralogs, although the *Tachyglossus* ancestral paralog appeared incomplete and was not used beyond confirmation of paralogous genes in egg-laying mammals. Placental and marsupial species each showed only a single copy of the derived paralog. These observations support a date for loss of the ancestral paralog in the lineage leading to therian mammals, rather than to all mammals. Gene models from several ray-finned fish (Actinopterygii) lineages support a third, lineage-restricted paralog with a more diverged protein sequence that has weaker match to the ANK motifs, GLYVGILK in place of the GLYLGYLK peptide, and no RA domain. However, neither *Danio* nor closely related *Danionella* models currently include a third paralog and the lineage-restricted paralogs from other species were not included in subsequent analyses.

We analyzed ANKFN1 protein sequence conservation with several tools. We first curated 208 homologous protein sequences from 159 diverse Holozoan organisms to represent full or near-full gene models across the well-conserved gene genealogy. Protein sequence models that performed poorly in multiple sequence alignment relative to sister groups and potential distant homologs in a Euglenoid and two Mucomycote fungi were excluded. We examined sequence conservation using Jensen-Shannon divergence [8] (Figure 1A). The most-conserved site in this analysis was the first tyrosine residue in the GLYLGYLK peptide (Figure 1B) and six of the eight residues in the peptide were among the most-conserved 2% of sites in the alignment. Similarly, a sequence-weighted composite score [9] (Figure 1C) placed three of the eight residues in the most-conserved 2% (Figure 1D). Multivariate analysis of protein polymorphism (MAPP) [10] was used to characterize physicochemical constraints based on amino acid representation and variability. Plotting the median substitution score (Figure 1E) again highlighted the GLYLGYLK sequence, with the consecutive GLY residues each in the top 2% of scores (Figure 1F) and two additional sites in the top 5%. This was the only sequence in the Holozoan ANKFN1 alignment with three consecutive positions in the top 5% of scores.

**Figure 1.**
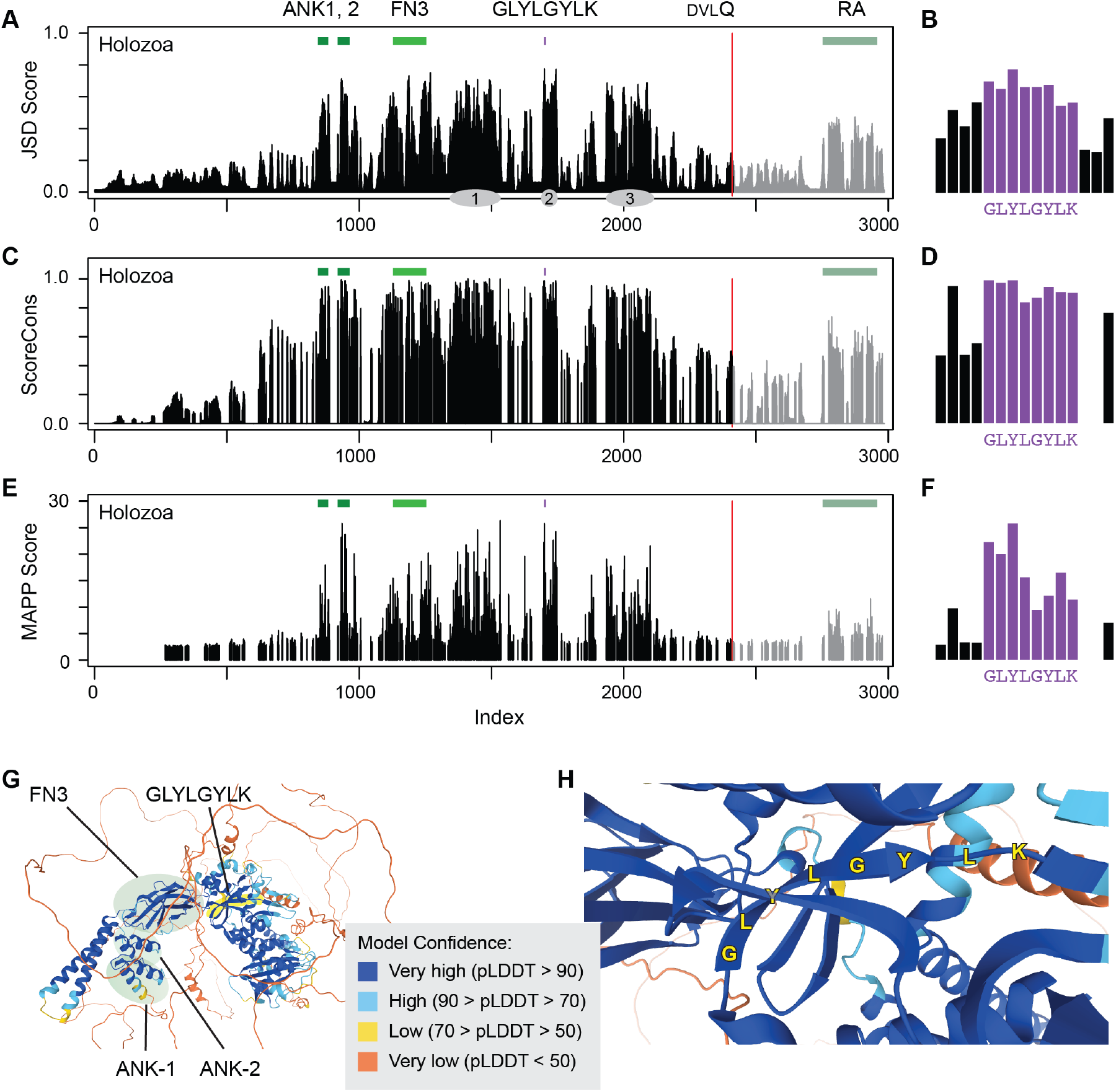
Conserved features of *Ankfn1* homologs. For each histogram (A-F) a higher bar indicates higher conservation or constraint at a single residue site for an alignment of Holozoan ANKFN1 proteins. (**A**) Jensen-Shannon divergence (JSD) scores standardized to a range of zero to one. Colored bars above the histogram indicate the positions of the two ankyrin repeats (ANK1,2) and fibronectin type 3 (FN3) domains, the conserved GLYLGYLK peptide, and the ras association (RA) domain of ancestral paralogs. A vertebrate-conserved DVLQ peptide (red line) marks the carboxyterminal extent of homology between ancestral and derived paralogs. Gray ovals indicate approximate positions of the three conserved non-motif regions of Zhang et al. [1]. (**B**) JSD scores at the conserved peptide, y-axis same as A. (**C**) ScoreCons residue conservation using Valdar’s scoring method. (**D**) ScoreCons scores at the conserved peptide, y-axis same as C. (**E**) Multivariate analysis of protein polymorphism (MAPP) median score for substitution at each site. (**F**) MAPP scores at the conserved peptide, y-axis same as E. (**G**) AlphaFold2 predicted structure for mouse ANKFN1 protein, showing positions of the ANK, FN, and GLYLGYLK (yellow highlight) regions and confidence scores. (**H**) Detail view of the structure in G showing the predicted GLYLGY beta strand in relation to flanking strands.

The predicted structure of human and mouse ANKFN1 proteins by AlphaFold2 [11, 12] is further instructive. The conserved GLYLGYLK peptide is predicted to reside in a compact structure of which it is one of the earlier (carboxyterminal) sequence elements (Figure 1G). The GLYLGY residues are predicted with high confidence to form a beta sheet internal to this structure and interacting with flanking beta sheets as structural components (Figure 1H). Disruption of this site might be expected to be highly deleterious to ANKFN1 function. Paralog-specific alignments from 49 species for which both were available showed strong conservation of the RA domain among ancestral homologs, but lower apparent conservation for C-terminal sequences among derived paralogs (Supplementary Figure S1). Consistent with both the high degree of conservation and predicted structural requirement, previous mutations at this sequence in both mice and flies were reported as presumptive null alleles [1].

### Simultaneous editing of both zebrafish *Ankfn1* homologs

We targeted the site encoding the GLYLGYLK peptide for mutagenesis of each paralog because it is highly constrained, resides in a frame-shifting exon, and the homologous site was previously targeted to create phenotypically null alleles in flies and apparent null mutations in mice [1], and should be similarly essential to the function of each zebrafish gene. We co-injected single-guide RNAs targeting the sequence encoding the GLYLGYLK peptide of each *D. rerio* paralog with Cas9 mRNA to allow simultaneous editing at both loci (Figure 1C). Injected G0 founders were outcrossed to wildtype fish and genomic DNA was prepared from G1 offspring to screen for germline-transmitted mutations. Among 60-90 injected offspring, we recovered a single frameshifting 11-bp deletion allele (*sd67*) on chromosome 12 and both an in-frame 18-bp deletion (*sd68*) and a frame-shifting 23-bp insertion allele on chromosome 24 (*sd69*). G1 fish from sequence-verified founders were raised to sexual maturity, genotyped from fin biopsies, and then intercrossed to produce animals for behavioral phenotyping [13].

### Highly penetrant major phenotypes require biallelic inactivation of both Ankfn1 homologs

Initial crosses during allele characterization demonstrated strong effects on swim bladder inflation, vestibular function, and survival to adulthood in animals that had both alleles of both *Ankfn1* homolog mutated (Tables 1-3). We assessed gross behavioral and physiological phenotypes in mutant offspring showed a minority of larvae with balance and locomotor deficits, followed by failure of swim bladder inflation. Larvae whose swim bladders failed to inflate died within two weeks and were not available for subsequent analysis.

**Table 1.** Cross: Chr12 (−11/+) Chr 24 (+23/+) x Chr12 (−11/+) Chr 24 (−18/+)

| Chr12 | Chr24 | Unaffected | Affected | Total |
| --- | --- | --- | --- | --- |
| -11/-11 | +23/-18 | 0 | 4 | 4 |
| -11/-11 | +23/+ | 9 | 0 | 9 |
| -11/-11 | -18/+ | 5 | 0 | 5 |
| -11/-11 | +/+ | 7 | 1 | 8 |
| -11/+ | +23/-18 | 7 | 0 | 7 |
| -11/+ | +23/+ | 7 | 0 | 7 |
| -11/+ | -18/+ | 14 | 0 | 14 |
| -11/+ | +/+ | 7 | 0 | 7 |
| +/+ | +23/-18 | 4 | 0 | 4 |
| +/+ | +23/+ | 8 | 0 | 8 |
| +/+ | -18/+ | 4 | 0 | 4 |
| +/+ | +/+ | 5 | 0 | 5 |
|  | Total | 77 | 5 | 82 |

**Table 2.** Cross: Chr12 (−11/+) Chr 24 (+23/+) x Chr12 (−11/-11) Chr 24 (+23/+)

| Chr12 | Chr24 | Unaffected | Affected | Total |
| --- | --- | --- | --- | --- |
| -11/-11 | +23/+23 | 0 | 10 | 10 |
| -11/-11 | +23/+ | 10 | 0 | 10 |
| -11/-11 | +/+ | 5 | 0 | 5 |
| -11/+ | +23/+23 | 6 | 0 | 6 |
| -11/+ | +23/+ | 14 | 0 | 14 |
| -11/+ | +/+ | 8 | 0 | 8 |
|  | Total | 43 | 10 | 53 |

**Table 3.**
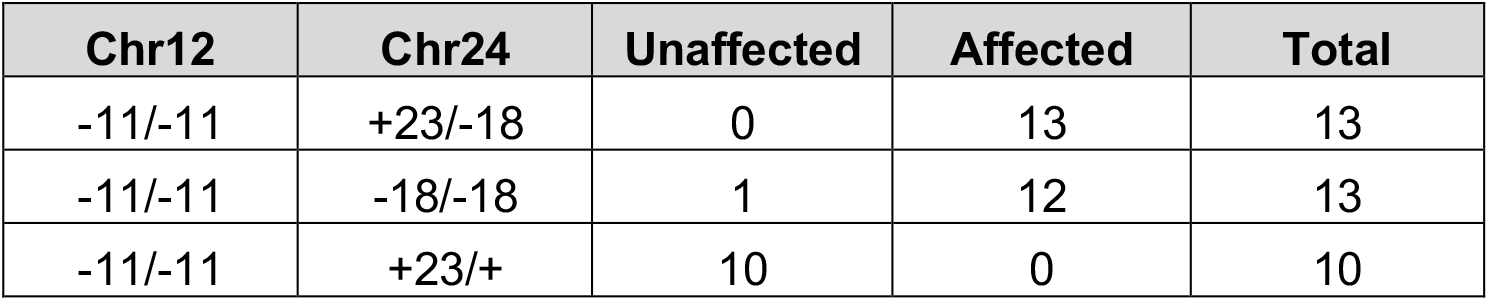
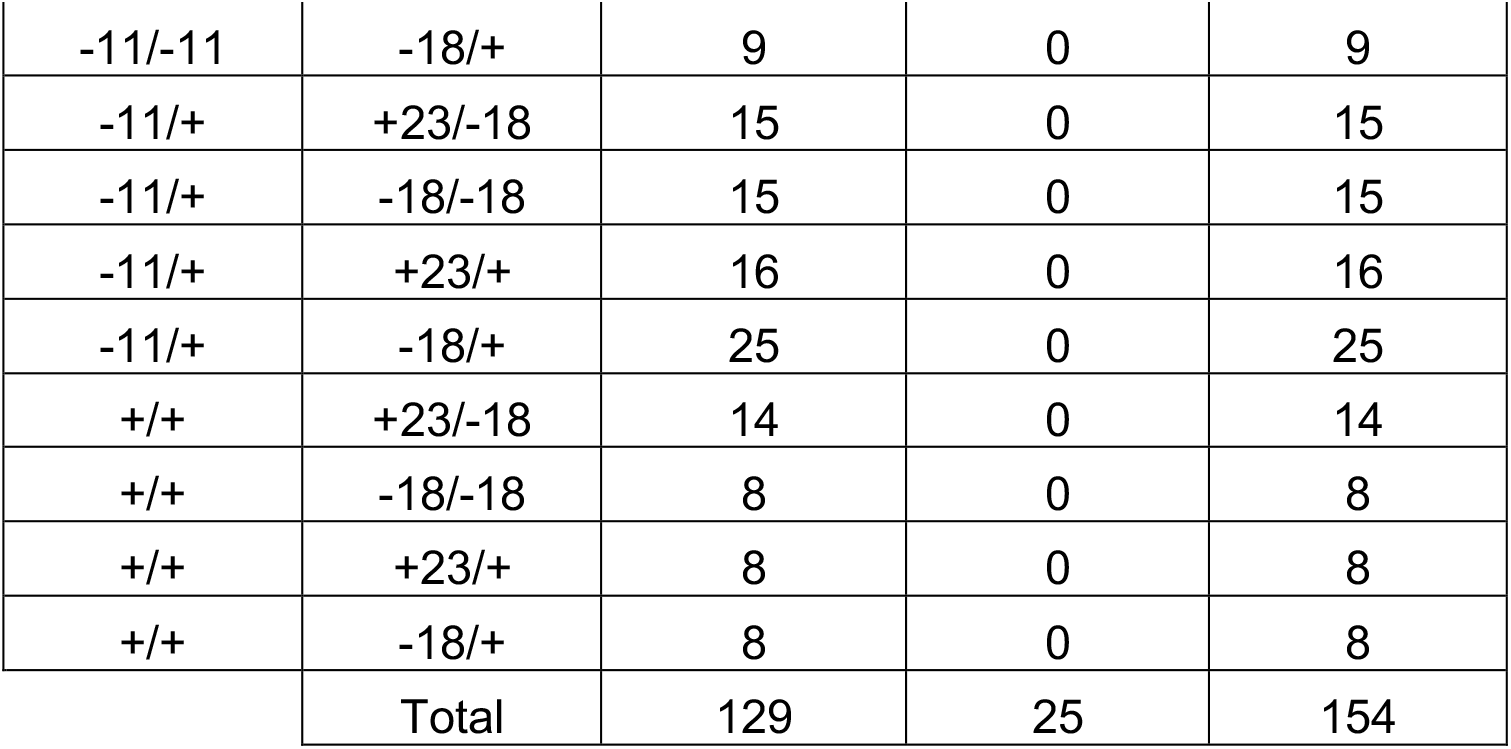
Cross: Chr12 (−11/+) Chr 24 (−18/+) x Chr12 (−11/+) Chr 24 (−18/+23)

We then set up formal dihybrid crosses of the frameshifting Chr12 and Chr24 mutant alleles in two cohorts (Table 4). Fish with both genes inactivated displayed abnormal balance, abnormal escape response to touch, and subsequent failure of swim bladder inflation (Figure 3, side view). Double mutant animals failed to maintain a dorsal-up posture (Figure 3, top view). Progeny were sorted categorically at swim bladder inflation and genotyped by PCR.

**Table 4.** Cross: Chr12 (−11/+) Chr 24 (+23/+) x Chr12 (−11/+) Chr 24 (+23/+)

| Chr12 | Chr24 | Unaffected | Affected | Total |
| --- | --- | --- | --- | --- |
| -11/-11 | +23/+23 | 1 | 15 | 16 |
| -11/-11 | +23/+ | 32 | 5 | 37 |
| -11/-11 | +/+ | 17 | 2 | 19 |
| -11/+ | +23/+23 | 29 | 0 | 29 |
| -11/+ | +23/+ | 56 | 1 | 57 |
| -11/+ | +/+ | 27 | 1 | 28 |
| +/+ | +23/+23 | 12 | 0 | 12 |
| +/+ | +23/+ | 35 | 2 | 37 |
| +/+ | +/+ | 19 | 0 | 19 |
|  | Total | 228 | 26 | 254 |

**Figure 2.**
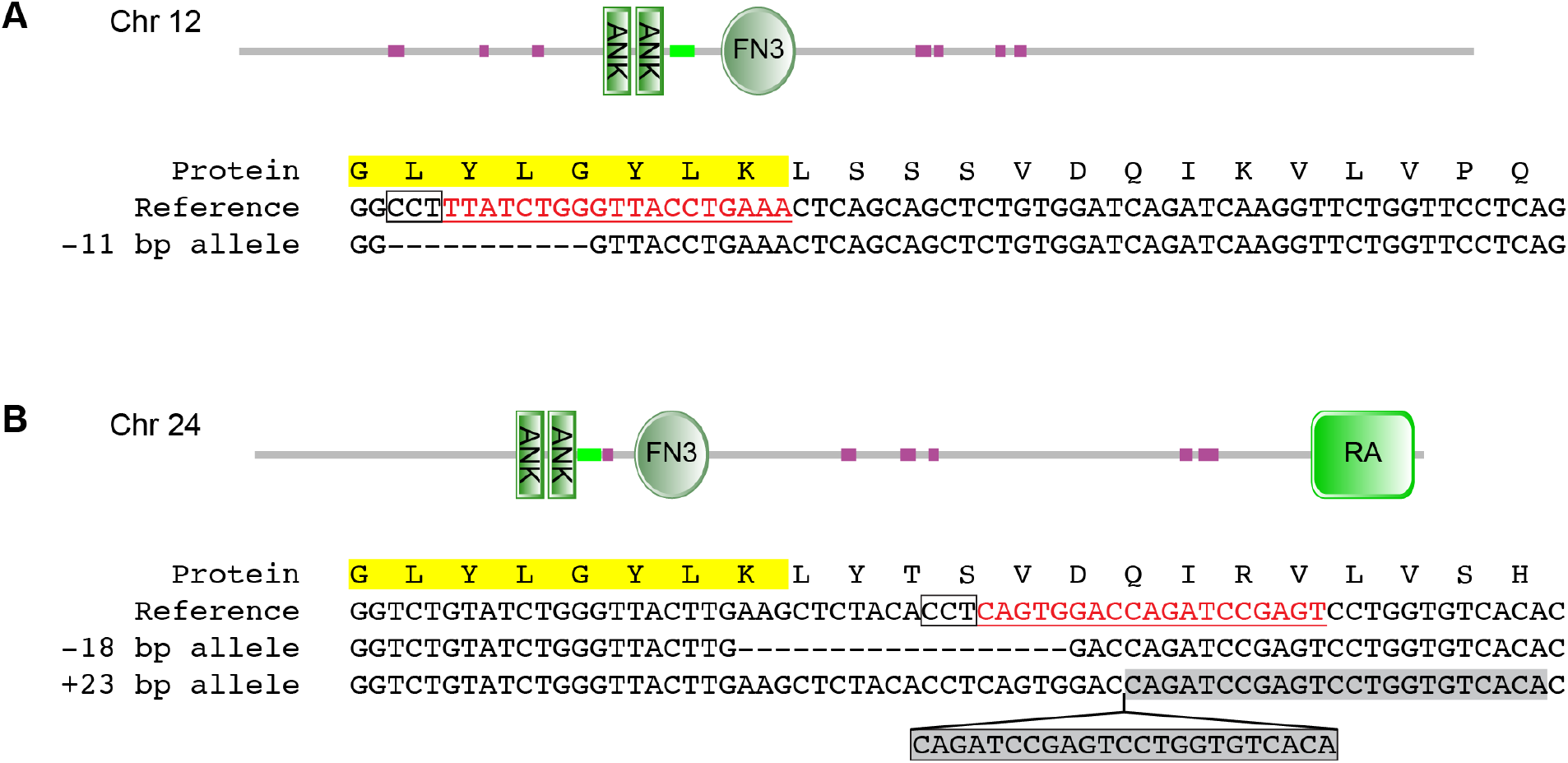
Targeted mutagenesis of *Ankfn1* homologs. Peptide and genomic sequences for targeted sites in *D. rerio Ankfn1* homologs. (**A**) *Ankfn1* on chromosome 12 is a derived paralog that includes two ANK and one FN3 domain but no RA domain. (**B**) *Ankfn1-like* on chromosome 24 is an ancestral paralog including the RA domain. The targeted GLYLGYLK site is highlighted in yellow. CRISPR guide RNA sites are underlined and protospacer adjacent motifs (PAMs) are boxed in each reference sequence. Recovered mutations are shown with deleted base pairs as dashes and insertion of a frame-shifting duplicated sequence (gray shadow) at chromosome 24 shown in a box below the caret.

**Figure 3.**
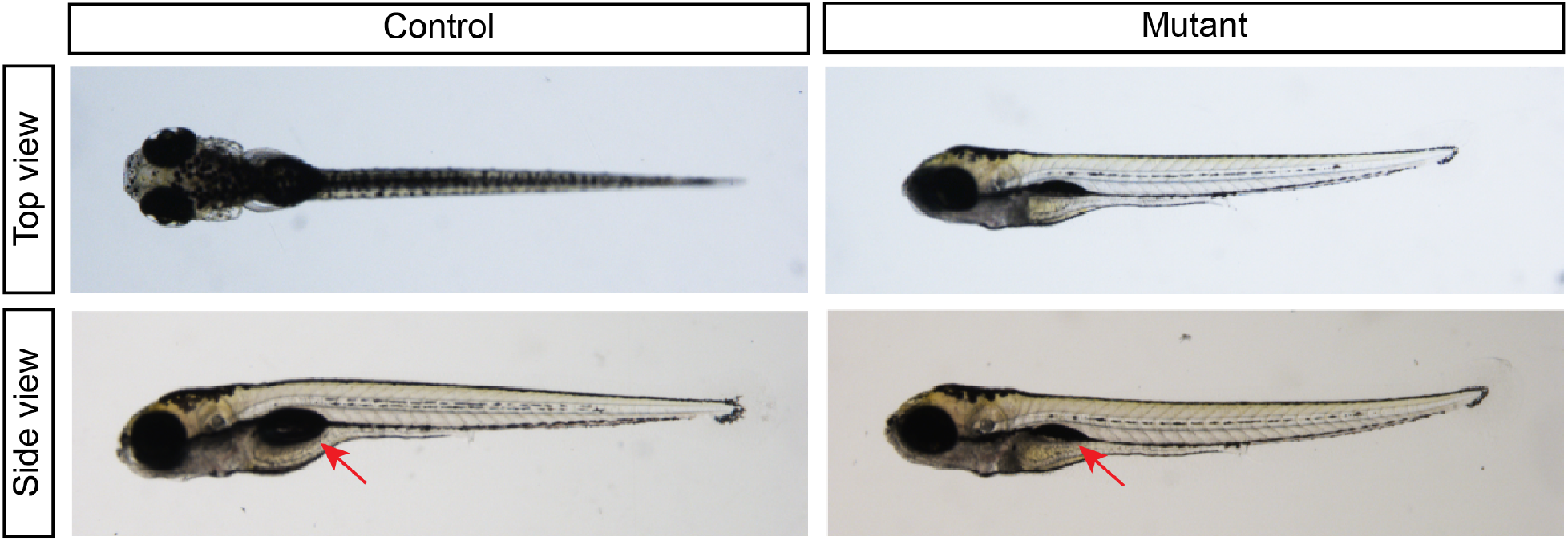
Swim bladder and posture defects in *Chr12/Chr24* mutant zebrafish. By five days post-fertilization mutant animals frequently failed to maintain dorsal orientation while swimming (Top view). Swim bladders typically failed to inflate in mutant animals (Side view, red arrows).

Genotype ratios of the offspring did not deviate from Mendelian expectations at 5 days post-fertilization (Chi square p=0.73), suggesting good viability to ascertainment. Among 254 offspring of the dihybrid crosses we observed that all double mutants, with one exception, failed to inflate their swim bladders within five days post-fertilization (94%). Comparatively, the frequency of inflation failure among all other genotypes was low, 11 out of 238 animals (4.6%). Interestingly, among the symptomatic animals that were not double mutants, the majority (7/11) where homozygous mutant for the Δ11 allele on chromosome 12, the derived copy which lacks the RA domain.

Pooling data from all crosses allowed an assessment of penetrance for each genotype class (Table 5). Categorical phenotypes in double mutant animals showed 96% (54/56) penetrance after combining results from both Chr24 mutations. For Chr12 homozygotes with at least one wild-type allele at Chr24 penetrance was 7% (8/112), including three affected animals that were wild-type at Chr24. Only four of the remaining 375 offspring had any observed phenotype and those four did not congregate by genotype class after correcting for sample sizes of the subgroups, with no affected offspring among 110 that had both alleles at Chr24 mutated but at least one wild-type allele at Chr12. This analysis support low penetrance of the Chr12 (derived) locus alone and strong synthetic interaction with the Chr24 (ancestral) locus for the behavioral and swim bladder inflation phenotypes.

**Table 5.** Summary of all crosses.

| Chr12 | Chr24 | Unaffected | Affected | Total | Penetrance |
| --- | --- | --- | --- | --- | --- |
| -11/-11 | +23/+23 | 1 | 25 | 26 | 0.96 |
| -11/-11 | +23/-18 | 0 | 17 | 17 | 1.00 |
| -11/-11 | -18/-18 | 1 | 12 | 13 | 0.92 |
| -11/-11 | +23/+ | 61 | 5 | 66 | 0.08 |
| -11/-11 | -18/+ | 14 | 0 | 14 | 0.00 |
| -11/-11 | +/+ | 29 | 3 | 32 | 0.09 |
| -11/+ | +23/+23 | 35 | 0 | 35 | 0.00 |
| -11/+ | +23/-18 | 22 | 0 | 22 | 0.00 |
| -11/+ | -18/-18 | 15 | 0 | 15 | 0.00 |
| -11/+ | +23/+ | 93 | 1 | 94 | 0.01 |
| -11/+ | -18/+ | 39 | 0 | 39 | 0.00 |
| -11/+ | +/+ | 42 | 1 | 43 | 0.02 |
| +/+ | +23/+23 | 12 | 0 | 12 | 0.00 |
| +/+ | +23/-18 | 18 | 0 | 18 | 0.00 |
| +/+ | -18/-18 | 8 | 0 | 8 | 0.00 |
| +/+ | +23/+ | 51 | 2 | 53 | 0.04 |
| +/+ | -18/+ | 12 | 0 | 12 | 0.00 |
| +/+ | +/+ | 24 | 0 | 24 | 0.00 |
|  | Total | 477 | 66 | 543 | 0.12 |

## Discussion

What happens to paralogous copies after gene duplication across broad time scales is an interesting general question, for which Ankfn1 is an interesting specific example. All invertebrate lineages examined have one copy, except urochordates, which have none. All vertebrate lineages have both an ancestral Ankfn1 homolog (including the RA domain) and a derived Ankfn1 homolog (lacking the RA domain), except therian mammals, which have lost the ancestral copy. The two-homolog arrangement has been conserved across non-therian lineages since it arose between the most recent common ancestor of jawed vertebrates ∼465 Mya and the most recent common ancestor of all vertebrates ∼615 Mya [14]. The single ancestral gene is essential in Drosophila, while the single derived gene impacts vestibular and neurological phenotypes in mice.

We used *D. rerio* as a model vertebrate for a first look at *Ankfn1* genetic properties in a vertebrate with paralogous copies. We directed mutations to the most highly conserved *Ankfn1* domain of each gene and recovered presumptive null frameshift alleles. By design, these mutations mirrored alleles previously reported for single homologs in mice and flies [1]. While evolution acts on a much finer scale than laboratory phenotypes and single locus deletions may have subtler or less-penetrant phenotypes than we had power to see, we found that biallelic inactivation of both Chr12 and Chr24 *Ankfn1* homologs was required to produce striking phenotypes with near complete penetrance. Inactivation of the Chr12 (derived) paralog alone was sufficient for weak penetrance of the larval and swim bladder phenotypes (7% of offspring), while inactivation of the Chr24 (ancestral) paralog had no detected effect on these phenotypes (0/110).

Our analysis supports functional overlap of *Ankfn1* paralogs in normal development of zebrafish vestibular function, consistent with the single derived copy in mice. We found a high penetrance for vestibular-related phenotype in animals homozygous for inactivating mutations in each paralog, but not in other genotype combinations. The present study is limited in that we did not perform detailed phenotyping of inner ear structures or perform expression level or localization measurements to determine whether disruption of one paralog resulted in increased expression of the other as a form of dosage compensation [15]. Whether more subtle phenotypes occur with each mutant singly will require further investigation. We had limited power to assess the potential for low-penetrance phenotypes in some genotype combinations.

Few examples of paralogous gene pairs in zebrafish for which disruption of both copies creates a phenotype that differs from knockout of either paralog alone have been published [16]. Our results provide a striking example, with a gross morphological phenotype arising from disruption of both members of a paralogous gene pair without an overt phenotype in single disruption of either member alone. By generating single and double mutations, we demonstrated that despite differences in their domain structure, there is substantial genetic redundancy between paralogous *Ankfn1* genes in zebrafish, despite loss of the RA domain in the derived copy and maintenance of independent paralogs across most lineages during roughly half a billion years of vertebrate evolution.

## Materials and Methods

### Sequence analysis

Previously reported *Ankfn1* homologs [1] were updated to more recent annotations and supplemented with new homologs identified through iterative BLASTP searches [17] targeting previously unavailable species. Only one species per genus was included unless species showed 10% or greater non-identity between orthologs (Drosophila species group). Gene models annotated as low-quality were excluded. Domain annotation for each included sequence was characterized using SMART [18, 19]. For non-mammal vertebrates, inclusion required each paralog to include their expected domain composition. Species, accession numbers, taxonomic groups, and annotated domains for each included sequence are provided in Supplemental Table S1. Curated homologs were aligned using MUSCLE [20] from the European Bioinformatics Institute web portal (https://ebi.ac.uk/Tools). Sequence conservation of aligned residues was evaluated using Valdar’s scoring method in ScoreCons [9] and using Jensen–Shannon divergence [8]. Physico-chemical and evolutionary constraints on specific positions were assessed using the median replacement score in MAPP [10].

### Genome editing

Mutant fish were generated by injecting a cocktail of sgRNAs and Cas9 mRNA into *Danio rerio* strain AB one-cell-stage embryos [21]. Ankfn1-homolog sgRNAs were in vitro transcribed from PCR-amplified templates (Supplemental Table S2). Synthetic Cas9 mRNA was transcribed from pCS2-nCas9n (gift of Dr. Wenbiao Chen, Addgene plasmid # 47929). The injection cocktail contained 200 ng/ul Chr12 sgRNA, 300 ng/ul Chr24 sgRNA, and 300 ng/ul Cas9 mRNA. Founder mutations were identified by pooling G1 embryos from individual G0 injected fish and screening for length polymorphisms by PCR. PCR products from putative mutants were Sanger sequenced to identify specific alleles. Husbandry and stock maintenance were carried out at 28°C as described [22, 23].

### Breeding and epistasis

G0 founders were outcrossed to wildtype fish to produce G1 offspring. Animals in cross 1 were G2 offspring from an intercross of G1 mutants. Crosses 2 and 3 were G4 and G5 fish from intercross matings. After phenotyping, single zebrafish larvae were harvested into individual wells in DNA extraction buffer for use in genotyping assays. 1 µl from a 50 µl suspension was used for each 20 µl PCR reaction. Primers for genotyping are given in Supplemental Table S3.

## Acknowledgements

We thank Danni Chen for fish husbandry and helpful comments. This work was supported in part by grants R01 GM086912 and R01 NS097534 to B.A.H. from the National Institutes of Health. K.D.R. was supported in part by Ruth L. Kirschstein Institutional National Research Service Award T32 GM008666 from the National Institute for General Medical Sciences.

---

**Figure S1.**
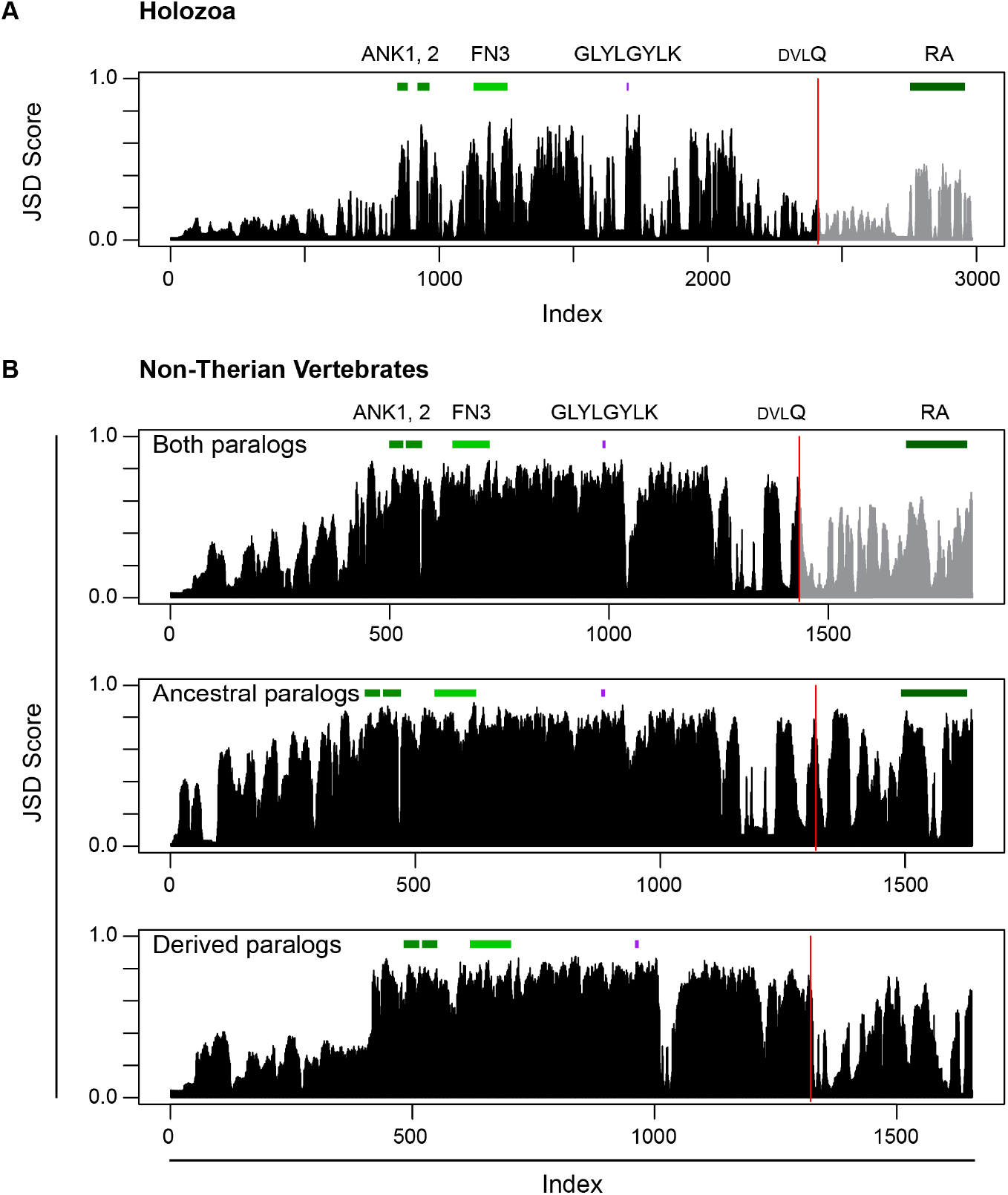
Paralog-specific conservation of vertebrate *Ankfn1* homologs. (**A**) Histogram of JSD scores for an alignment of Holozoan homologs with positions of annotated features repeated from Figure 1A for comparison. (**B**) JSD scores for aligned sites among 98 ANKFN1 protein homologs from 49 distinct species of non-therian jawed vertebrates for which both paralogs passed nominal filters for completeness showed higher scores and more compact alignment than the full Holozoa set due to reduced sequence diversity. Paralogs diverge after the DVLQ peptide (red line), indicated by gray histogram after this point in the joint alignment. Separate alignment of ancestral paralogs showed conservation within the RA domain broadly consistent with other domains. Derived paralogs showed less conservation after the DVLQ point among derived paralogs.

## Notes

### Competing Interest Statement

The authors have declared no competing interest.

